# Cortico-spinal Mechanisms of TENS-induced Analgesia: A Single-Blind, Three-Arm, fMRI-Based Randomized Controlled Trial

**DOI:** 10.1101/2024.02.06.579059

**Authors:** Zhaoxing Wei, Yunyun Duan, Yupu Zhu, Xiaomin Lin, Leiyao Zhang, Ming Zhang, Jonathan C.W. Brooks, Yaou Liu, Li Hu, Yazhuo Kong

## Abstract

**Background:** Transcutaneous Electrical Nerve Stimulation (TENS) is widely used for pain management, yet how different parameters—Conventional (high frequency, low intensity) versus Acupuncture-Like (low frequency, high intensity)—modulate pain remains controversial. These modes produce variable analgesic effects, but their underlying brain–spinal mechanisms are unclear. Clarifying these pathways could refine TENS protocols, optimize clinical outcomes, and reduce opioid reliance, highlighting the need for more precise, non-pharmacological approaches to advanced pain care.

**Methods:** This single-blind, three-arm randomized trial recruited 95 healthy adults (18–30 years) and assigned them (1:1:1) to Conventional TENS, Acupuncture-Like TENS, or sham. Participants underwent a 30-minute TENS intervention on the left forearm (C5–C6 dermatome) with simultaneous brain–spinal fMRI before and after thermal nociceptive stimuli. The primary outcomes were changes in Numeric Rating Scale (NRS) pain scores and brain–spinal BOLD signals. Secondary outcomes included psycho-physiological interaction (PPI) and mediation analyses of periaqueductal gray (PAG) activity and brain– cord connectivity.

**Findings:** Both TENS modes significantly reduced pain but engaged distinct cortico-spinal pathways. Conventional TENS yielded local analgesia via dlPAG-driven spinal inhibition plus partial cortical involvement (PAG–vmPFC). By contrast, Acupuncture-Like TENS produced diffuse analgesia through vlPAG-linked top-down modulation reliant on spinal gating. Correlation and mediation analyses confirmed that Conventional TENS integrates spinal and cortical synergies, whereas Acupuncture-Like TENS is dominated by robust descending control.

**Interpretation:** Different TENS parameters yield distinct analgesic mechanisms. Conventional TENS couples direct spinal inhibition with partial cortical regulation, while Acupuncture-Like TENS relies heavily on top-down pathways passing through the spinal cord. Recognizing these unique descending networks can guide targeted TENS protocols for diverse pain conditions, optimizing clinical outcomes and reducing reliance on pharmacological approaches.

**Fundings:** This work was supported by the National Key R&D Program of China (2022YFC3500603), the National Natural Science Foundation of China (32071061, 82072010, 82030121 and 32100861, 82330057), the Beijing Natural Science Foundation (IS23108, JQ22018).

## Introduction

Pain is a multifaceted experience affecting large populations worldwide, often culminating in reduced quality of life and imposing a substantial socioeconomic burden (1). The opioid crisis has brought increased attention to the side effects of pharmacological analgesic methods. Nonpharmacological approaches, neuromodulation in particular, are emerging as preferred pain treatments. Non-invasive neuromodulatory techniques, such as Transcutaneous Electrical Nerve Stimulation (TENS) that target the peripheral nervous system, do not necessitate surgical procedures (2).

TENS employs alternating currents with varying frequencies and intensities to stimulate peripheral nerves (3). Known for being safe, convenient, and low cost, TENS has effectively mitigated various pain conditions, e.g., postoperative pain and low back pain (3). TENS generally offers two primary modes: Conventional TENS generates analgesia within targeted dermatomes with higher frequency (∼50-100 Hz) and lower intensity (evoking a nonpainful sensation); Acupuncture-Like TENS produces a diffuse analgesic effect that extends beyond the dermatome, with lower frequency (∼2-4 Hz) and higher intensity (elicits a tolerable of painful sensation) (4, 5).

Previous studies revealed the analgesic effect of non-invasive neuromodulation, but provided limited evidence of the precise neurobiological mechanisms, particularly regarding how they interact with the complex networks of the central nervous system (6). Research using animal models has suggested that TENS relieves pain by activating peripheral nerve fibers, thereby modulating (inhibitory or excitatory) pain processing in the central nervous system, along the spinal cord, periaqueductal gray (PAG) of the brainstem, and pain-related cortical regions (7). Specifically, Conventional TENS modifies GABA levels in the dorsal horn and stimulates *δ*-opioid receptors in the spinal cord and brainstem, while Acupuncture-Like TENS primarily targets local *μ*-opioid receptors (8). Two theories predominantly explain the efficacy of TENS. Firstly, Conventional TENS is believed to achieve pain relief primarily through spinal inhibition (9), in line with the gate control theory and influenced by supraspinal areas (10). Secondly, both TENS modes activate systemic mechanisms, such as the endogenous opioid system or diffuse noxious inhibitory controls (DNIC) (3, 11), affecting the cortex, brainstem, and notably, the spinal cord.

Recent human studies employing electroencephalography (EEG) and functional magnetic resonance imaging (fMRI) have demonstrated that TENS can modulate spontaneous brainstem activity, particularly in the pons, subnucleus reticularis dorsalis (SRD) and rostral ventromedial medulla (RVM) (5). Moreover, Conventional TENS has been shown to enhance functional connectivity between the PAG and lateral prefrontal cortex (PFC) (10). Acupuncture-Like TENS induces more significant changes in alpha oscillations within the primary sensorimotor cortex (S1/M1) and increases the functional connectivity between S1 and the medial PFC more than Conventional TENS (4). However, human studies directly investigating the entire cortico-brainstem-spinal circuit mechanism in neuromodulation are lacking, leading to a gap in the comprehensive understanding of non-invasive neuromodulation analgesia in human research.

Over the past decade, simultaneous cortico-spinal fMRI has become pivotal in human pain research. This method provides an exceptional opportunity to capture neural activities across the cortex, brainstem, and spinal cord, thus providing systemic insights into the complex neural mechanisms involved in pain processing and neuromodulation (12). Within this framework, the functional networks in the cortico-brainstem-spinal pathway, mainly centering around the PAG, play a multifaceted role in pain modulation. This is evidenced in various contexts, including nocebo hyperalgesia (13), opioid analgesia (14), and attentional analgesia (15). Consequently, our current study aims to investigate the analgesic mechanisms of both Conventional and Acupuncture-Like TENS, focusing on the comprehensive cortico-brainstem-spinal pathway, especially PAG related endogenous analgesia pathway. Building upon existing literature, we hypothesize that both TENS modes engage the cortico-spinal pain modulation system; their distinct analgesic effects may derive from different neural interactions within this pathway, with the PAG as a relay site but potentially involving direct spinal inhibition or diffuse noxious inhibitory controls.

## Methods

### Study design and Participants

This study was a single-blind, three-arm, parallel-group randomized controlled trial conducted at Beijing Tiantan Hospital, China. Ninety-five healthy adults (aged 18–30 years, all right-handed, with no history of chronic pain, mental illness, or analgesic use, and no prior TENS treatment) were invited to participate. Due to equipment failure, three participants were unable to complete the study. Thus, the final analysis included ninety-two participants (50 females, age (mean ± SD) = 21.9 ± 3.23). The study was conducted in accordance with the Declaration of Helsinki and received approval from the Ethics Committee of the Institute of Psychology at the Chinese Academy of Science (H19023). Participants received a compensation of 200 RMB for their participation.

### Randomisation and Masking

Participants were randomly allocated (1:1:1) to one of three groups: Conventional TENS (*N*=30), Acupuncture-Like TENS (*N*=31), or sham TENS (*N*=31), each receiving a specific TENS treatment. The randomization sequence was generated by an independent statistician using computer-generated random numbers. The TENS operator was unblinded for proper parameter adjustment, while participants remained blinded. To minimize placebo effects and expectation biases, participants were not informed that the study pertained to analgesia, and electrical stimulation was not described as a therapeutic intervention. Instead, all participants were instructed to sit quietly, focus on the stimulation site, and were told, “An electrical stimulation will be continuously applied to your left forearm for the next 30 minutes. Please close your eyes and feel this stimulation.”

### Procedure and Interventions

Specifically, our experiment adopted a three-way mixed design with a 3 (group: Conventional, Acupuncture-Like, sham) × 2 (side: left treated side, TS; right untreated side, UTS) × 2 (order: pre-TENS, post-TENS) paradigm (Figure 1A to C).

**Figure 1.**
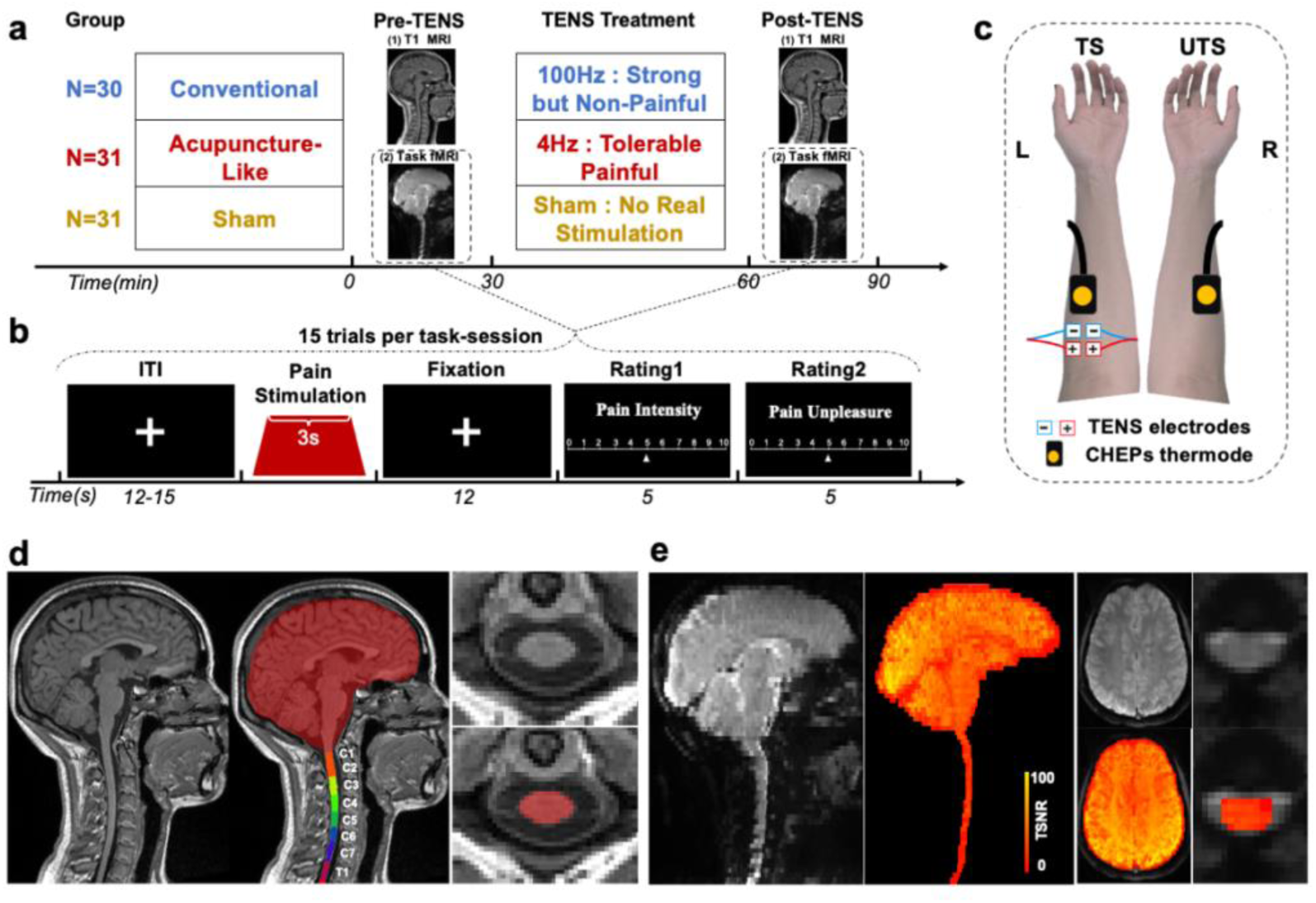
Experimental paradigm. (A) Overview of the experimental process. Participants were assigned to one of three experimental groups to receive different TENS treatments. Cortico-spinal images were acquired before and after TENS treatment. (B) Heat pain fMRI paradigm. Heat pain stimulation was applied for 3 seconds, followed by pain intensity and unpleasantness ratings. 15 trials were conducted for each run. (C) TENS treatment and heat pain stimulation. TENS electrodes were applied to the C5∼C6 dermatome of the left side, while the CHEPS thermode was applied to the same dermatome of both the left and right sides. The left is the treated side (TS), and the right is the untreated side (UTS). (D) Bias-corrected T1 images and segment brain and spinal cord examples from one representative participant. (E) Data quality check: distortion and physiological noise corrected fMRI image and corresponding temporal signal-to-noise ratio (tSNR) map of segmented cortico-spinal areas (sagittal and transversal views) from one representative participant.

The TENS treatment was delivered using a multichannel electrical stimulator (SXC-4A, Beijing Sanxia Technology Co., Ltd., China)(5). Two pairs of electrodes (diameter: 16 mm; interelectrode distance: 3 cm) were attached to the C5∼C6 dermatomes based on Natter’s Atlas of Human Neuroscience (16) in the ventrolateral part of the left forearm (treatment side, TS) (Figure 1C). The electrical stimuli in TENS consisted of a series of constant-current square-wave pulses, with the TENS treatment parameters varying across experimental groups and each stimulation lasting 30 minutes (4, 5) (Figure 1A). The Conventional TENS group received high-frequency stimulation at 100 Hz, with a 2000 µs pulse width and intensity set at 6.04 ± 1.44 mA, inducing a strong but nonpainful tingling sensation without causing muscle contractions. The Acupuncture-Like TENS group received low frequency stimulation at 4 Hz, with a 4000 µs pulse width and higher intensity of 7.00 ± 1.42 mA, set to elicit severe but tolerable pain and muscle twitching, akin to acupuncture needle sensation. The sham TENS group was informed of receiving faint electrical stimulation, but no actual current was delivered.

During the pre- and post-TENS sessions, Heat pain stimulation was applied to the same dermatome (C5∼C6) on both the left (treated) and right (untreated) sides using a Pathway device with a CHEPS thermode (Medoc Ltd, Ramat Yishai, Israel). Pain intensity and unpleasantness were rated on a Numeric Rating Scale (NRS, 0–10) following each stimulus, and the order of hands was pseudorandomized across participants. Simultaneous cortico-spinal scans of structural and task-based functional MRI with heat stimuli were obtained before and after TENS treatment.

### Statistic analysis

Behavioral data statistical analyses were performed using *jamovi* 1.2 (The *jamovi* project, 2020, https://www.jamovi.org). Imaging data were processed with FSL 6.0.6.5 (17) (FMRIB software library) and SCT 5.6 (18) (Spinal Cord Toolbox). The mean pain intensity and unpleasantness ratings for each block were calculated for each participant. A 3 (group: Conventional, Acupuncture-Like, Sham) × 2 (side: TS; UTS) × 2 (order: pre-TENS, post-TENS) mixed-factorial ANOVA was used to estimate the main effect and interactions of behavioral data. Post hoc within-group 2 “side” × 2 “order” repeated measures ANOVA, paired t-tests, and correlation analyses were subsequently performed for the behavioral data and brain-spinal fMRI data. P < 0.05 was considered as statistically significant for all behavioral analyses. Whole brain analyses were performed with a cluster-wise correction (*Z* > 3.1, *P* < 0.05). The brainstem and spinal analyses were subjected to TFCE correction (*P* < 0.05, threshold-free cluster enhancement) after 10,000 permutations. We performed a psycho-physiological interaction (PPI) analysis with the PAG as a seed to explore the functional interaction along the cortico-brainstem-spinal endogenous analgesia pathway. In the following step, we constructed a GLM mediation analysis using the ’jAMM’ package in the *jamovi* to inspect the potential mediation effect of Δconnectivity and analgesic effect (Δpain intensity and Δunpleasantness) across each group.

The primary outcomes were changes in subjective pain ratings (NRS) and alterations in brain–spinal BOLD signals. Secondary outcomes included PPI and mediation analyses focusing on PAG connectivity. Additional details regarding questionnaires, quantitative sensory testing (QST), MRI data acquisition and preprocessing, and advanced statistical analyses are provided in the Supplementary Materials.

## Results

### Psychophysics of pain intensity and unpleasantness with TENS modulations

One-way analysis of variance (ANOVA) revealed no significant differences among the three groups in demographic variables, pain cognition, and emotion (Table S1), as well as Quantitative Sensory Test (QST) indices (Table S2). A 3 “group” × 2 “side” × 2 “order” mixed-design ANOVA revealed significant interaction effects for pain intensity (*F*(_2,89_) = 28.80, *P* < 0.001) and unpleasantness (*F*(_2,89_) = 14.50, *P* < 0.001), indicating different analgesic effects across the three TENS groups.

To further dissect the analgesic effects, we implemented a 2 (side: TS, UTS) × 2 (order: pre-TENS, post-TENS) repeated-measures ANOVA within each TENS treatment group (Figure 2A and D). In the Conventional TENS group, significant “side” × “order” interaction effects emerged for pain intensity (*F*_(1,29)_ = 38.60, *P* < 0.001) and unpleasantness (*F*_(1,29)_ = 20.70, *P* < 0.001). Post hoc paired t-tests revealed a significant analgesic effect in TS (pain intensity: t_(29)_ = 6.87, P < 0.001, unpleasantness: *t*_(29)_ = 5.25, *P* < 0.001) but not in UTS, suggesting a localized analgesic effect of Conventional TENS within the treated dermatome. In the Acupuncture-Like TENS group, we observed significant main effects for “order” (pain intensity: *F*_(1,30)_ = 37.94, *P* < 0.001; unpleasantness: *F*_(1,30)_ = 19.13, *P* < 0.001) and “side” (pain intensity: *F*_(1,30)_ = 21.50, *P* < 0.001; unpleasantness: *F*_(1,30)_ = 11.17, *P* = 0.002), but no significant interaction effect. Post hoc paired t-test found a significant analgesic effect in both the TS (pain intensity: *t*_(30)_ = 5.47, *P* < 0.001; unpleasantness: *t*_(30)_ = 4.47, *P* < 0.001) and UTS (pain intensity: *t*_(30)_ = 6.04, *P* < 0.001; unpleasantness: *t*_(30)_ = 3.75, *P* < 0.001), indicating diffuse analgesic effects even for untreated dermatome from Acupuncture-Like TENS. The sham TENS group had no significant analgesic effect in either TS or UTS.

**Figure 2.**
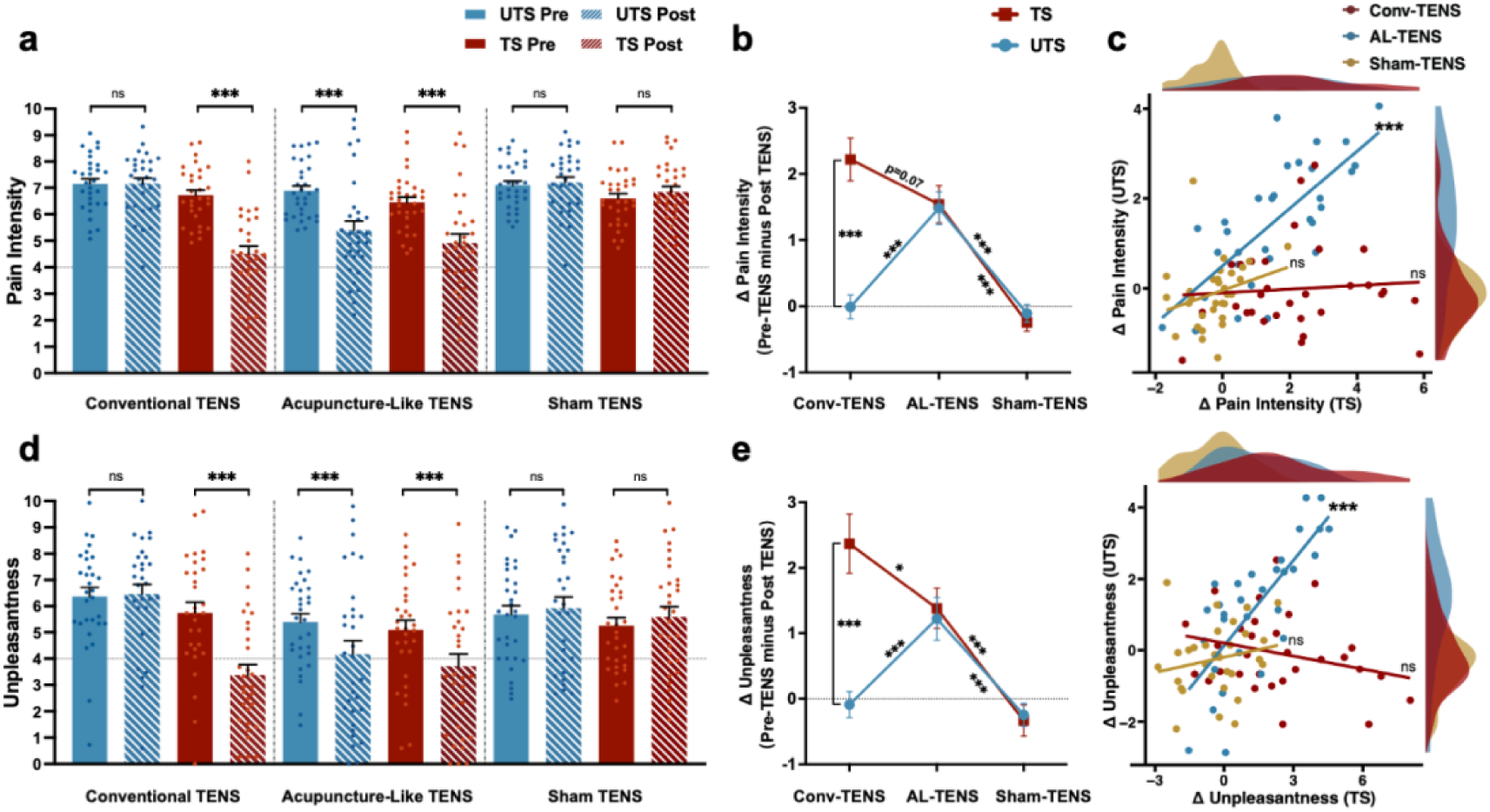
Behavioral ratings. (A and D) Pain intensity and unpleasantness rating results of the three groups. In the Conventional TENS group, pain perception was significantly reduced after treatment only in TS. In the Acupuncture-Like TENS group, both the TS and UTS demonstrated significant reductions in pain perception. However, neither side showed significant pain modulation in the sham TENS group. (B and E) Analgesic effect (pre minus post) of pain intensity and unpleasantness showed the strongest effect in the TS of the Conventional TENS group. (C and F) Correlations of the analgesic effects (Δpain intensity and Δunpleasantness) between TS and UTS in each group revealed a significant positive correlation only for the Acupuncture-Like TENS group. Dots represent the individual participant values, and error bars indicate the SEM. * *P* < 0.05; ** *P* < 0.01;*** *P* < 0.001; ns: not significant.

To directly quantify and compare the analgesic effects (Δpain intensity and Δunpleasantness, pre-TENS minus post-TENS) across groups, we conducted a 3 (group: Conventional, Acupuncture-Like, Sham) × 2 (side: TS, UTS) mixed-design ANOVA (Figure 2B and E). We detected a significant interaction effect between “group” and “side” in Δpain intensity (*F*_(2,89)_ = 27.10, *P* < 0.001) and Δunpleasantness (*F*_(2,89)_ = 14.50, *P* < 0.001), suggesting different analgesic effects among the three TENS modes. The Conventional TENS group exhibited significant differences between TS and UTS in terms of Δpain intensity (*t*_(89)_ = - 8.79, *P* < 0.001) and Δunpleasantness (*t*_(89)_ = -6.64, *P* < 0.001). However, no differences were observed between TS and UTS in the Acupuncture-Like and sham TENS groups.

Additionally, we employed Pearson correlation analysis to investigate the relationship between TS and UTS analgesic effects within each group (Figure 2C and F). A significant positive correlation between TS and UTS analgesic effects (Δpain intensity: *r* = 0.73, *P* < 0.001; Δunpleasantness: *r* = 0.76, *P* < 0.001) was observed exclusively in the Acupuncture-Like TENS group, i.e. regardless of the treatment location, both sides showed consistent analgesic pattern. No such correlation was detected in the Conventional TENS or sham TENS group.

### Brain and spinal cord blood-oxygen-level-dependent (BOLD) activations

Group average BOLD activations from the cervical cord to the cortex, induced by heat pain stimulation prior to TENS treatment, were observed in classical pain-associated regions (Figure S2). These included primary and secondary somatosensory cortices (S1 and S2), primary motor cortex (M1), supplementary motor area (SMA), anterior cingulate cortex (ACC), insula, thalamus, cerebellum, PAG, RVM (cluster-forming threshold *Z* > 3.1, family-wise error (FWE) corrected *P* < 0.05), and the dorsal areas of C6 segments of the spinal cord (*P* < 0.05, threshold-free cluster enhancement (TFCE) corrected).

To investigate TENS-induced modulation of activities across the entire cortico-spinal pathway, we implemented a group-level General Linear Model (GLM) model of 2 (side: TS, UTS) × 2 (order: pre-TENS, post-TENS) repeated-measures ANOVA on the simultaneous cortico-spinal fMRI data for each group. Please note that to control the lateralization bias between left- and right-side nociceptive stimulation (19) we flipped the X dimension of each subject’s right-side heat pain-activated first-level parameter estimation images for higher-level group comparisons. Whole brain analysis after multiple comparisons indicated that Conventional TENS exerted a localized modulatory effect (interaction) on heat pain-evoked activation in the TS but not in UTS at the cortico-spinal level (Figure 3). Conversely, Acupuncture-Like TENS demonstrated a diffuse modulatory effect (main effect of “order”) on heat pain-evoked activation in both the TS and UTS (Figure 4).

**Figure 3.**
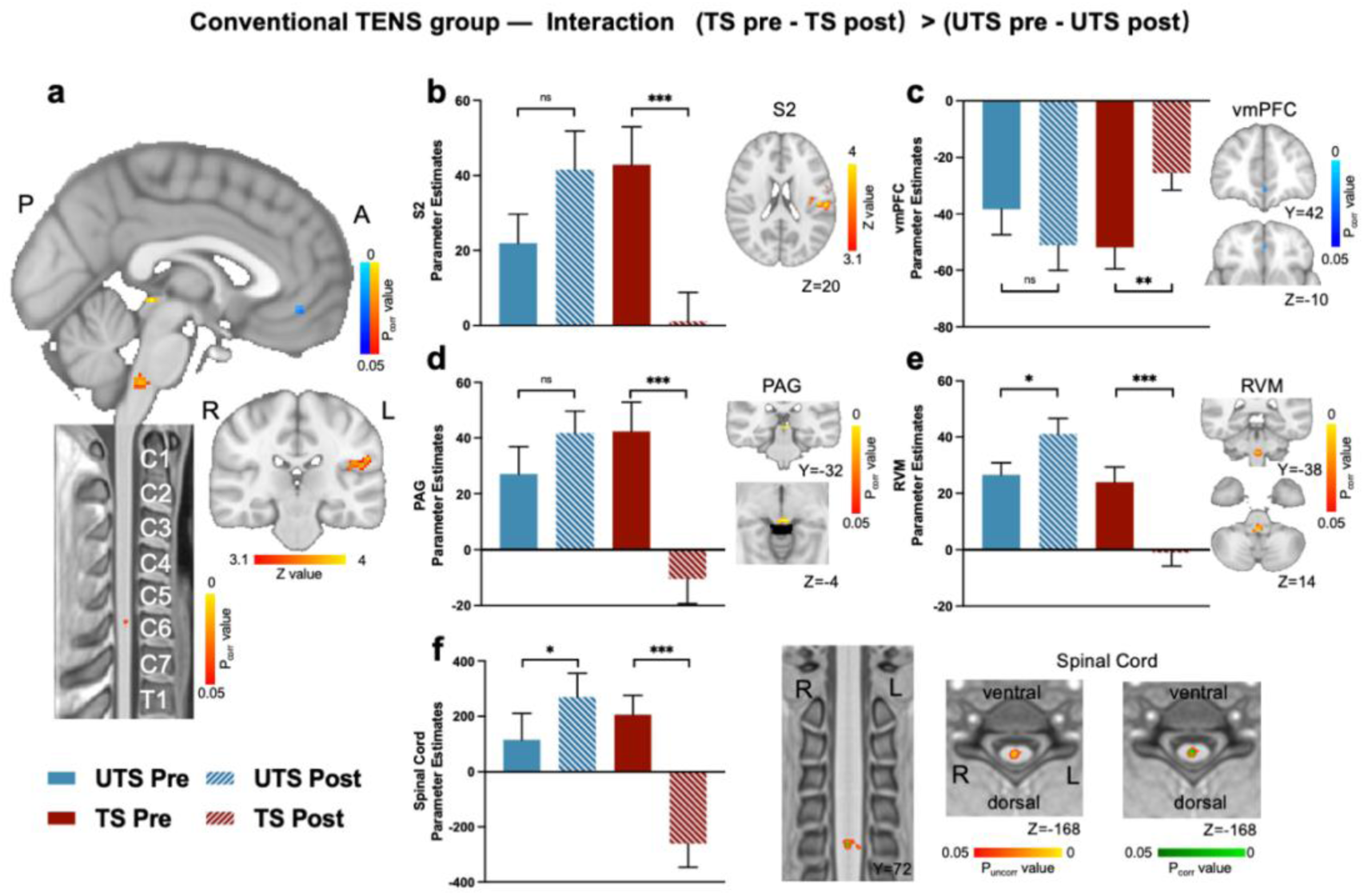
Modulations of BOLD responses in the brain and spinal cord due to Conventional TENS. (A) The Conventional TENS group showed a significant interaction effect in the left S2 (cluster forming threshold *Z* > 3.1, FWE corrected *P* < 0.05), vmPFC, PAG, RVM, and spinal cord C6 segment (*P* < 0.05, TFCE corrected). (B) Parameter estimates (PE) of ipsilateral S2 showed a significant decrease for TS after TENS treatment. (C) PEs of vmPFC showed a significant increase for TS after TENS treatment. (D) PEs of dorsal lateral PAG showed a significant decrease of TS after TENS treatment. (E) PEs of RVM showed a significant decrease for TS and an increase for UTS after TENS treatment. (F) The first panel: PEs of the C6 spinal cord showed a significant decrease for TS after TENS treatment; the second panel: coronal view of statistical *z*-maps overlaid on a PAM50 T1 image; the third panel: axial view with the threshold set to *P*_uncorr_ < 0.05 for visualization (red); the fourth panel: axial view with the threshold set to *P*_corr_ < 0.05. * *P* < 0.05; ** *P* < 0.01; *** *P* < 0.001; ns: not significant.

**Figure 4.**
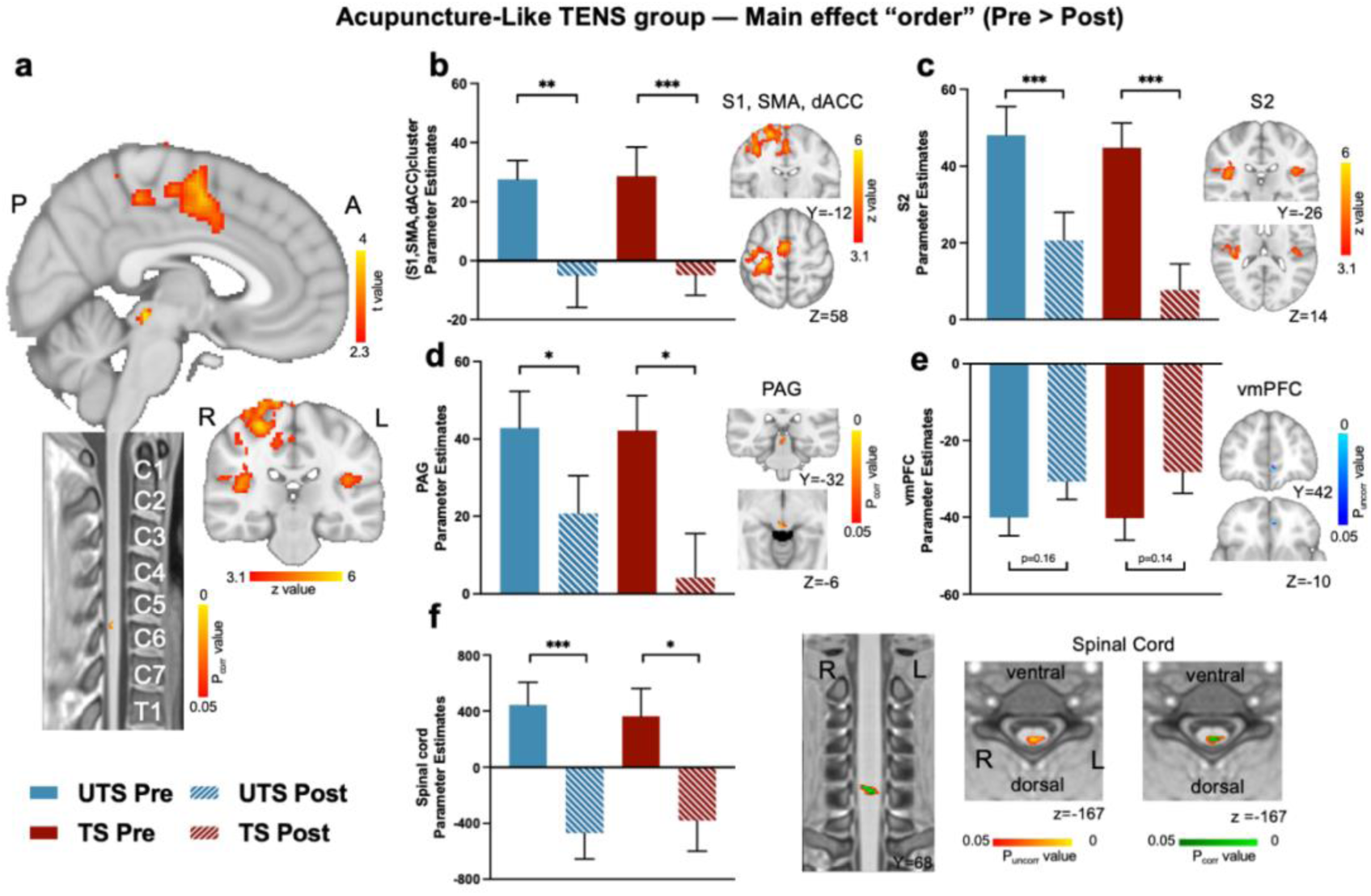
Modulation of BOLD responses in the brain and spinal cord due to Acupuncture-Like TENS. (A) The Acupuncture-Like TENS group showed a significant main effect of “order” in the S1, S2, dACC, SMA (cluster forming threshold *Z* > 3.1, FWE corrected *P* < 0.05), PAG, and spinal cord C6 segment (*P* < 0.05, TFCE corrected). (B) PEs of right S1, dACC, SMA cluster showed decreases in both TS and UTS after TENS treatment. (C) PEs of bilateral S2 showed decreases in both TS and UTS after TENS treatment. (D) PEs of ventral lateral PAG showed decreases in both TS and UTS after TENS treatment. (E) PEs of vmPFC showed non-significant increases in both TS and UTS after TENS treatment. (F) The first panel: PEs of the spinal cord showed decreases in both TS and UTS after TENS treatment; the second panel: coronal view of statistical z-maps overlaid on a PAM50 T1 image; the third panel: axial view with the threshold set to *P*_uncorr_ < 0.05 for visualization (red ‒ yellow); the fourth panel: axial view with the threshold set to P_corr_ < 0.05 (green). * *P* < 0.05; ** *P* < 0.01;*** *P* < 0.001.

Specifically, within the Conventional TENS group, a significant interaction contrast showed several pain-associated regions along the cortico-spinal pathway (Figure 3), including cortical areas of the left S2 (cluster forming threshold *Z* > 3.1, FWE corrected *P* < 0.05), ventromedial PFC (vmPFC), PAG, RVM as well as spinal cord C6 segments (TFCE corrected, *P* < 0.05). Parameter estimates (PEs) were extracted from the regions showing significant interaction effects for post-hoc t-tests. In particular, the S2 (*t*_(29)_ = 4.03, *P* < 0.001), PAG (*t*_(29)_ = 4.78, *P* < 0.001), RVM (*t*_(29)_= 3.73, *P* < 0.001), and spinal cord (*t*_(29)_ = 4.59, *P* < 0.001) demonstrated significant decreases in heat pain activation, while the vmPFC exhibited a significant increase in activation (*t*_(29)_ = -3.33, *P* = 0.002) in TS. In the UTS, we detected a non-significant modulation in activation in the S2 and PAG, while significant activation increases were observed in RVM (*t*_(29)_ = -2.61, *P* = 0.01) and spinal cord (*t*_(29)_ = -2.23, *P* = 0.03). No significant decrease in vmPFC activation (*t*_(29)_ = 1.41, *P* = 0.17) was found in response to heat pain stimulation in the UTS (Figure 3B to F).

We did not identify regions with significant interaction effects within the Acupuncture-Like TENS group. However, we observed the significant main effect of “order” in several regions, including the right SMA, dorsal ACC (dACC), bilateral S1, S2, left cerebellum (cluster-forming threshold *Z* > 3.1, FWE corrected *P* < 0.05), PAG, as well as spinal cord C6 segment (*P* < 0.05, TFCE corrected, Figure 4). Parameter estimates were extracted for post-hoc t-tests from regions with a significant main effect of “order”. ROI analysis indicated significant activity decreases in both TS and UTS within the S1, SMA, dACC cluster (TS: *t*_(30)_ = 3.64, *P* = 0.001; UTS: *t*_(30)_ = 2.93, *P* = 0.006), bilateral S2 (TS: *t*_(30)_ = 5.42, *P* < 0.001; UTS: *t*_(30)_ = 4.62, *P* < 0.001), PAG (TS: *t*_(30)_ = 2.70, *P* = 0.01; UTS: *t*_(30)_ = 2.01, *P* = 0.05) and spinal cord (TS: *t*_(30)_ = 2.40, *P* = 0.02; UTS: *t*_(30)_ = 3.62, *P* = 0.001). Unlike the Conventional TENS group, the vmPFC did not show a significant change of activities in either TS (*t*_(30)_ = -1.53, *P* = 0.14) or UTS (*t*_(30)_ = -1.43, *P* = 0.16) (Figure 4B to F).

The sham TENS group did not exhibit either significant main effect of “order” or interaction effect. Other detailed brain ANOVA results for each group are provided in Figure S3.

### PAG activity associated with local and diffuse analgesic effects of TENS

Our study specifically focused on the PAG, a core region involved in pain perception and the descending modulation of pain signals. We examined the relationship between ΔPAG activity (pre-TENS minus post-TENS) and the analgesic effects (Δpain intensity, also pre-TENS minus post-TENS) in both TENS groups. In the Conventional TENS group, ΔPAG activity significantly positively correlated with the Δpain intensity (*R²* = 0.45, *β* = 0.01, *SE* = 0.00, *t* = 2.55, *P* = 0.01), and the difference in effects between the TS and the UTS was statistically significant (*β* = -1.71, *SE* = 0.41, *t* = -4.20, *P* < 0.001). Further correlation analysis revealed a significant positive correlation between Δpain intensity and ΔPAG activity in the TS (*r* = 0.39, *P* = 0.04) but not in the UTS (*r* = 0.22, *P* = 0.23) (Figure 5A and C). In the Acupuncture-Like TENS group, ΔPAG activity significantly positively correlated with the Δpain intensity (*R²* = 0.18, *β* = 0.01, *SE* = 0.00, *t* = 3.60, *P* < 0.001). However, the effect difference between the TS and the UTS was insignificant (*β* = 0.08, *SE* = 0.00, *t* = 3.60, *P* = 0.81) (Figure 5B and D).

**Figure 5.**
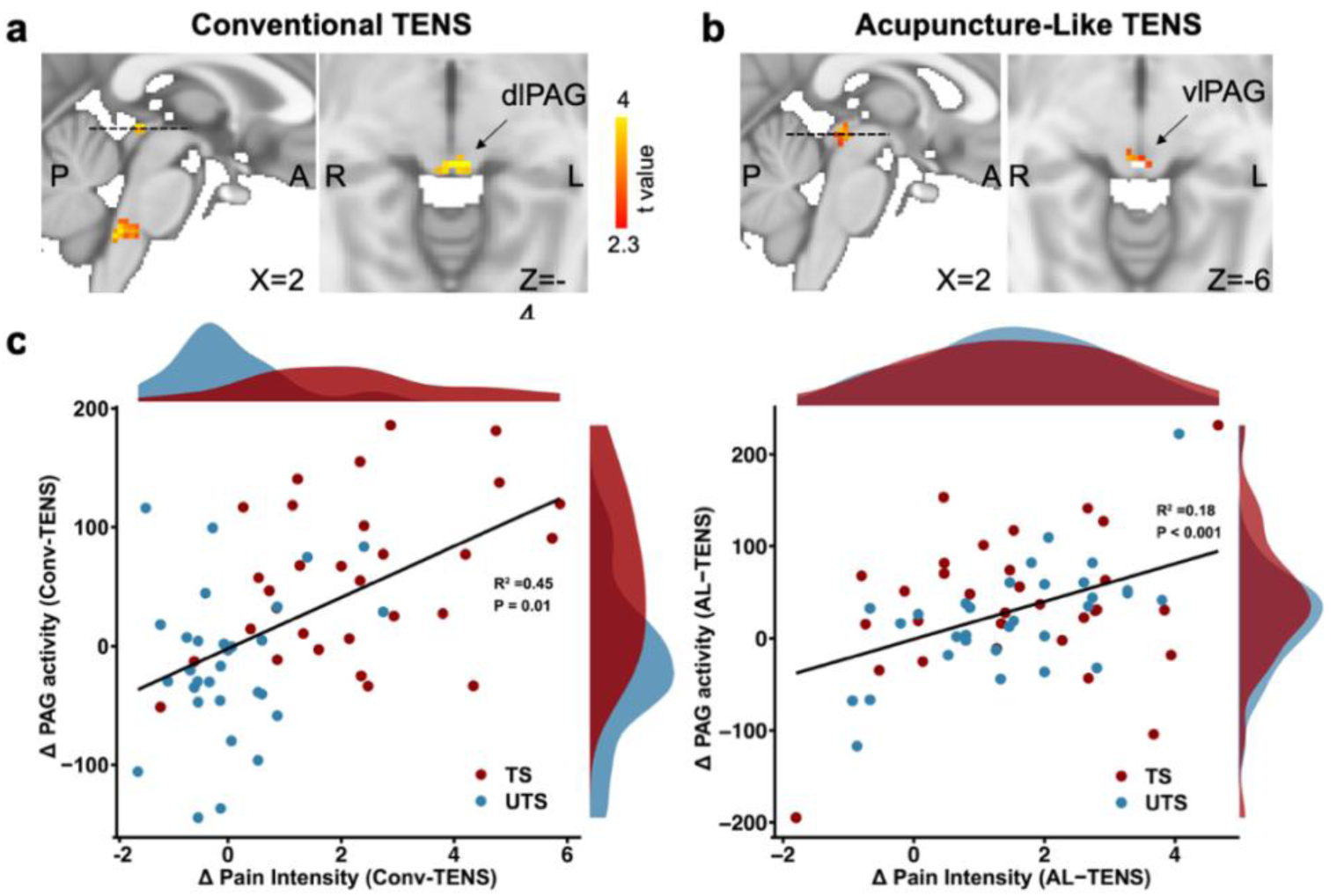
PAG activity correlated with the analgesic effect of TENS. (A) Left dlPAG (ipsilateral TENS treatment) significantly decreased TS activation in the Conventional TENS group. (B) Right vlPAG (contralateral TENS treatment) showed the main “order” effect in the Acupuncture-Like TENS group, in which both the TS and UTS showed a significant decrease in activation. (C) Conventional TENS group: Δdorsal lateral PAG (dlPAG) activity has a significant positive correlation with the Δpain intensity (*R²* = 0.45, *β* = 0.01, *P* = 0.01), and the difference in effects between the TS and the UTS was statistically significant (*β* = -1.71, *P* < 0.001). (D) Acupuncture-like TENS group: Δventral lateral PAG (vlPAG) activity has a significant positive correlation with the Δpain intensity (*R²* = 0.18, *β* = 0.01, *P* < 0.001). However, the effect difference between the TS and the UTS was insignificant (*β* = 0.08, *P* = 0.81).

### PAG-based cortical-spinal TENS analgesic pathways

To delve deeper into the cooperation between brain and spinal cord regions under different neuromodulation conditions along the entire cortico-spinal pathways, psycho-physiological interaction (PPI) analyses were employed. For the first-level PPI analysis, the PAG served as a seed region to explore its coupling with areas identified from ANOVA results and previous literature (4, 10), including bilateral S1, vmPFC, and C5-C7 cervical cord. A 2 × 2 within-group repeated-measures ANOVA was performed for the group-level PPI analysis. Subsequently, mediation analysis with 1000 bootstraps was applied to integrate significant PPI changes (Δconnectivity, pre-TENS minus post-TENS) with behavioral analgesic effects to uncover the potential mechanisms behind TENS-induced analgesia. We find different contributions of PAG-spinal and PAG-cortical connectivities in mediating the analgesic effects elicited by the two modes of TENS.

In the Conventional TENS group (Figure 6A), a significant interaction effect was noted for PAG-vmPFC connectivity (left vmPFC, ipsilateral to TENS stimulus, *P* < 0.05, TFCE corrected), primarily dominated by a significant increase in TS (*t*_(29)_ = -2.10, *P* = 0.04) and a non-significant decrease in UTS. For PAG-spinal connectivity, a significant interaction effect (*P* < 0.05, TFCE corrected) was also found due to a significant decrease in TS (*t*_(29)_ = 4.92, *P* <0.001) and a non-significant increase in UTS. Furthermore, the change in PAG-spinal connectivity (ΔPAG-spinal) was found to have a significant positive correlation with the Δpain intensity (*r* = 0.68, *P* < 0.001) and Δunpleasantness (*r* = 0.61, *P* < 0.001), but the change in PAG-vmPFC connectivity (ΔPAG-vmPFC) had a negative correlation with the Δpain intensity (*r* = -0.54, *P* < 0.001) and Δunpleasantness (*r* = -0.47, *P* < 0.001) (Figure 6B and Figure S5A). These results suggest that more substantial analgesic effect are paralleled by the more pronounced changes in these neural pathways. The mediation analysis revealed that ΔPAG-vmPFC connectivity partially mediates the relationship between ΔPAG-spinal connectivity and the analgesic effect. This was evident in both the Δpain intensity (indirect path = 0.01, *P* = 0.02, 95% C.I.= [0.004, 0.02], Figure 6E) and Δunpleasantness (indirect path = 0.01, *P* = 0.05, 95% C.I.= [1.39e-4, 0.02], Figure S5C). These findings suggest that following Conventional TENS treatment, PAG-spinal connectivity not only directly impacts pain perception but also exerts an indirect influence by modulating PAG-vmPFC connectivity.

**Figure 6.**
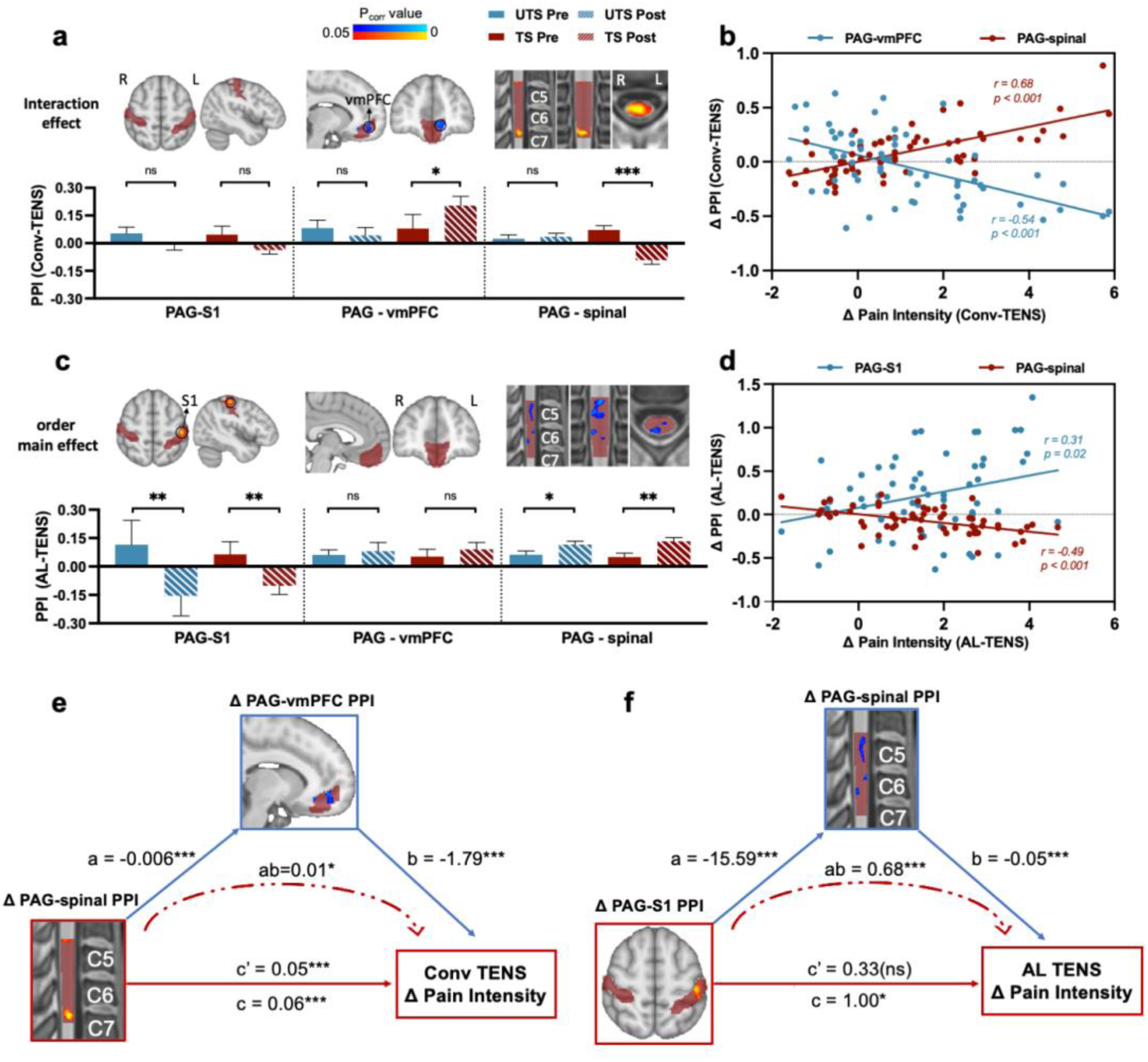
PAG-cortical and PAG-spinal connectivity modulate TENS analgesic effects. (A) PAG-vmPFC and PAG-spinal connectivity exhibited a significant interaction in the Conventional TENS group (*P* < 0.05, TFCE corrected), with only TS showing significant differences. (B) Conventional TENS analgesic effect (Δpain intensity) positively correlated with ΔPAG-spinal connectivity (*r* = 0.68, *P* < 0.001) and negatively correlated with ΔPAG-vmPFC connectivity (*r* = -0.54, *P* < 0.001). (C) PAG-S1 and PAG-spinal connectivity demonstrated a main effect of "order" in the Acupuncture-Like TENS group (*P* < 0.05, TFCE corrected). (D) Acupuncture-like TENS analgesic effect (Δpain intensity) positively correlated with ΔPAG-S1 connectivity (*r* = 0.31, *P* = 0.02) and negatively correlated with ΔPAG-spinal connectivity (*r* = -0.49, *P* < 0.001). (E) In the Conventional TENS group, ΔPAG-vmPFC connectivity partially mediated the effect of ΔPAG-spinal connectivity on Δpain intensity (indirect path = 0.01, *P* = 0.02, 95% C.I.= [0.004, 0.02]). (F) In the Acupuncture-Like TENS group, ΔPAG-spinal connectivity fully mediated the effect of ΔPAG-S1 connectivity on Δpain intensity (indirect path = 0.68, *P* = 0.007, 95% C.I.= [0.26, 1.27]). Note: ΔPAG-spinal connectivity values were normalized by dividing each value by 100 for visualization. * *P* < 0.05; ** *P* < 0.01; *** *P* < 0.001; ns: not significant.

In the Acupuncture-Like TENS group (Figure 6C), a significant "order" effect was found for the PAG-S1 (left S1, ipsilateral to TENS stimulus) and PAG-spinal connectivity (*P* < 0.05, TFCE corrected), resulting from a significant decrease in PAG-S1 and increase in PAG-spinal for both TS (S1: *t*_(30)_ = 2.88, *P* = 0.007; spinal: *t*_(30)_ = -2.94, *P* = 0.006) and UTS (S1: *t*_(30)_ = 2.78, *P* = 0.009; spinal: *t*_(30)_ = -2.20, *P* =0.04). The changes in PAG-S1 connectivity (ΔPAG-S1) had a significant positive correlation with Δpain intensity (*r* = 0.31, *P* = 0.02) and Δunpleasantness (*r* = 0.29, *P* = 0.02), and the change in PAG-spinal connectivity (ΔPAG-spinal) had a negative correlation with Δpain intensity (*r* = -0.49, *P* < 0.001) and Δunpleasantness (*r* = -0.41, *P* < 0.001) (Figure 6D and Figure S5B). These results also suggest that substantial analgesic effect are associated with pronounced changes in cortico-spinal pathways, however the PAG based associations seemed to be exactly opposite patterns for two TENS modes (Figure 6C vs Figure 6D). Further, the mediation analysis revealed that ΔPAG-spinal connectivity fully mediates the relationship between ΔPAG-S1 connectivity and the analgesic effect. This was evident in both the Δpain intensity (indirect path = 0.68, *P* = 0.007, 95% C.I.= [0.26, 1.27], Figure 6F) and Δunpleasantness (indirect path = 0.65, *P* = 0.02, 95% C.I.= [0.09, 1.20], Figure S5D). These findings suggest that following Acupuncture-Like TENS treatment, PAG-S1 connectivity can only indirect influence pain perception by modulating PAG-spinal connectivity.

## Discussion

To the best of our knowledge, this is the first study to utilize simultaneous cortico-spinal fMRI to investigate the central analgesic mechanism of a noninvasive periphery neuromodulation technique — TENS. Through a three-way mixed-design combining the latest cortico-spinal fMRI with heat pain stimuli (Figure 1A to C), we found that Conventional TENS primarily induces local analgesia effects on the treated side (Figure 2 and 3), while Acupuncture-Like TENS produces diffuse analgesic effects on both sides (Figure 2 and 4). A crucial discovery is the role of the PAG in TENS-induced analgesia, with activities in the dlPAG linked to Conventional TENS and vlPAG associated with Acupuncture-Like TENS (Figure 5). Importantly, direct spinal inhibition (PAG-spinal connectivity) partially mediated by PAG-vmPFC connectivity leads to local analgesic effects in Conventional TENS. In contrast, a top-down diffuse noxious inhibition (PAG-S1 connectivity) fully mediated by PAG-spinal connectivity leads to diffusion analgesic effects in Acupuncture-Like TENS (Figure 6). Our findings provide systematic neural evidence and a comprehensive understanding of the analgesic mechanisms induced by TENS.

These results were obtained by combining several recent advancements in simultaneous cortico-spinal fMRI to address the inherent difficulties in imaging a large area: simultaneous multi-slice imaging (SMS) with parallel image reconstruction to get shorter scan duration and less artifact aliasing; novel EPI distortion correction using reverse phase encoded B0s; robust physiological noise removal to increase tSNR and eliminate false positives; optimized analysis pipeline for separated brain and spinal cord images including preprocessing, co-registration, and group statistics, etc.

Understanding the analgesic mechanism of TENS is crucial for clinical applications of peripheral-based non-invasive neuromodulation. Two well-known and fundamental theories might be involved: gate control (9) and DNIC (20). Gate control theory suggests that activating Aβ sensory afferents with tactile stimuli can inhibit nociceptive signals at the spinal level. In contrast, the DNIC theory posits that nociceptive stimuli can suppress pain by activating the descending pain modulation system (20), altering signal input in the spinal cord. These theories involving systemic mechanisms encompass the cortex, subcortical regions, and the spinal cord. Our study observed distinct disparities in pain modulation between Conventional TENS and Acupuncture-Like TENS, evident in both behavioral outcomes (Figure 2) and cortico-brainstem-spinal BOLD responses and couplings (Figure 3 and 4), consistent with findings from animal studies (21).

Conventional TENS mainly provides pain relief limited to the directly treated area, as seen by a significant reduction of pain at the TS as opposed to the UTS (Figure 2A and D). Corresponding to these behavioral observations, Conventional TENS was found to significantly modulate activity in several regions activated by heat pain on the TS, specifically in areas such as S2, vmPFC, PAG, RVM, and the spinal cord (Figure 3 and Figure S2). These findings robustly demonstrate the involvement of supraspinal levels in Conventional TENS analgesia. Specifically, vmPFC, which is involved in the emotional and cognitive aspects of pain and its modulation (22), was influenced by Conventional TENS. Additionally, PAG and RVM, crucial brainstem nuclei in the pain modulation system, also play a key role in this process. Recent human studies have shown that Conventional TENS can increase functional connectivity between PAG and lateral PFC (10), and increase cortico-spinal excitability, which is indicated by motor evoked potentials (MEP) elicited by TMS (11). Our results (Figure 3) support previous studies by showing that Conventional TENS works by blocking opioid receptors in the RVM and activating opioid receptors in the PAG (23).

Acupuncture-Like TENS demonstrated a more diffuse analgesic effect in behavioral outcomes (Figure 2A and D), which was evidenced by significant pain reduction observed on both the TS and UTS, with a notable positive correlation between the two sides (Figure 2C and F). Regarding BOLD responses, Acupuncture-Like TENS altered activations in a widespread range of regions evoked by heat pain stimulation on both sides. These regions include the right SMA, dACC, bilateral S1 and S2, left cerebellum, PAG, and spinal cord (Figure 4). A previous EEG study also revealed that Acupuncture-Like TENS could induce the spontaneous activity state changes within the sensory-motor cortex (S1/M1), and increase functional connectivity between S1 and the medial PFC (4). Furthermore, other research has shown that after Acupuncture-Like TENS treatment in carpal tunnel syndrome patients, there was a significant decrease in the activation intensities of S2, M1, bilateral SMA, and other regions compared to baseline (24). Recent studies on the cortico-spinal mechanisms of conditioned pain modulation (CPM) also demonstrate the importance of S2 (25).

An interesting finding is the differential S2 responses to TENS modes: Conventional TENS specifically reduced ipsilateral S2 pain activation (Figure 3B); in contrast, Acupuncture-Like TENS led to a bilateral decrease in S2 pain activation (Figure 4C). S2 encodes the intensity and location of pain, but is also involved in higher-order functions, including sensorimotor integration, synthesis of bilateral information, attention, learning, and memory (26). Animal studies have demonstrated that unilateral tactile stimulus could modulate neuronal activity in somatosensory cortices of the ipsilateral hemisphere (27), which may support Conventional TENS as a form of tactile stimulation, reducing ipsilateral S2 activation. Bilateral S2 is involved in pain processing across both humans and animals, the decrease in bilateral S2 activation further substantiates the widespread analgesic effect induced by Acupuncture-Like TENS.

The response patterns across various brain regions under different TENS modes may depend on different descending pain modulation pathways (14). Both TENS engage the PAG in their analgesic processes (Figure 3D, 4D and Figure S4B). Furthermore, The dlPAG is more actively involved in Conventional TENS, while the vlPAG is more prominent in Acupuncture-Like TENS (Figure 5). The PAG receives and transmits cortical inputs to the RVM and other key regions in the descending pain modulation system. This complex role is highlighted in various studies, including those on opioid analgesia (14), placebo analgesia (28), attention-driven analgesia (15), and expectations of pain response (29), all indicating intricate interactions within a broad network. Moreover, distinct subregions within the PAG contribute differently to pain modulation. Preclinical studies have shown that opioid-mediated analgesia involves neurons in the vlPAG, while non-opioid analgesia is associated with neurons in the lPAG and dlPAG. This differentiation suggests that the analgesic effect of Conventional TENS might be due to spinal-driven inhibition rather than opioid-mediated descending modulation. Conversely, Acupuncture-Like TENS’s efficacy seems to be linked to opioid-related descending modulation.

Focusing solely on the activations of brain and spinal cord regions seems insufficient; taking advantage of innovative cortico-spinal imaging, we further examine functional interactions along the central nervous system (30). The functional connectivity between the PAG-cortical and PAG-spinal induced analgesic effects provides crucial insights into the PAG’s role in pain modulation. We find that PAG-vmPFC connectivity showed significant alterations following Conventional TENS application (Figure 6A), and these changes correlated with analgesic effects (Figure 6B). Enhanced coupling between the PFC and PAG has been observed in placebo analgesia ; furthermore, a more substantial placebo analgesic effect is associated with increased coupling strength (31). Previous animal studies have demonstrated that subpopulations of mPFC neurons project to the PAG, specifically the dlPAG (32). In the Acupuncture-Like TENS group, changes in PAG-S1 were correlated with analgesic outcomes (Figure 6C and B). Existing rs-fMRI connectivity between the vlPAG and S1 is related to top-down pain modulation, and the vlPAG has efferent connections with the spinal cord, which is also related to opioid-mediated analgesia (33).

Additionally, significant alterations in PAG-spinal connectivity were observed following both TENS treatments, with these changes correlating with analgesic effects (Figure 6B and D). However, the direction of these changes differed between modes: in the Conventional TENS group, connectivity shifted from positive to negative, whereas in the Acupuncture-Like TENS group, positive connectivity was enhanced post-treatment (Figure 6A and C). Neural projections between the PAG and spinal cord are well-established, with their connectivity strength linked to pain intensity (34). Positive connectivity, where activation in one area boosts activity in another, indicates synergy, while negative connectivity suggests inhibition, as activation in one region diminishes activity in another. In the Conventional TENS group, reversing positive to negative PAG-spinal connectivity may signify spinal-driven inhibition, reducing pain signal transmission. Conversely, enhanced positive PAG-spinal connectivity in the Acupuncture-Like TENS group points to a systemic pain modulation mechanism, possibly opioid-mediated descending modulation, where increased connectivity enhances pain inhibitory pathways.

Our mediation results further illustrate that Conventional TENS-induced analgesia through both direct spinal inhibition (PAG-spinal connectivity) and descending modulation pathways (PAG-vmPFC connectivity) (Figure 6E). In contrast, Acupuncture-Like TENS’s diffusion analgesic effects appear to be fully induced by descending DNIC (PAG-S1 connectivity fully mediated by PAG-spinal connectivity) (Figure 6F). This further explicates that the strongest localized (i.e. treated side) behavioral analgesic effects induced by Conventional TENS (Figure 2B and E) result from the synergistic action of dual pathways, with the descending modulation pathway also eliciting non-significant changes in pain-induced perceptions and cortico-spinal activations in the UTS. Conversely, the analgesic effects elicited by Acupuncture-Like TENS exhibit diffuse and consistent characteristics across both subjective perceptions and cortico-spinal activations and connectivity, serving as compelling evidence of the DNIC’s effectiveness. These observations provide crucial insights into the neurobiological mechanisms underpinning the efficacy of these two TENS modes.

This study used an effective behavioral paradigm and simultaneous cortico-spinal imaging technology to systematically investigate the analgesic effects and cortico-brainstem-spinal mechanisms of TENS, a typical peripheral neuromodulation method. We observed differences in the analgesic effects of the two TENS modes at the behavioral level and confirmed distinct mechanisms by examining PAG-vmPFC, PAG-S1, and PAG-spinal connections. Most importantly, we demonstrated the crucial role of spinal inhibition and descending pain modulation in Conventional TENS analgesia and descending pain modulation in Acupuncture-Like TENS analgesia. Our results support TENS for pain management and open avenues for exploring cortico-spinal mechanisms in other neuromodulation techniques like TMS, tDCS, and spinal cord stimulation (SCS). However, this study has limitations, primarily using young healthy subjects for experimental acute pain, without utilizing chronic pain models (e.g., capsaicin) in a larger sample and across a wider age range, or involving TENS analgesia in actual patients. Future research will delve deeper into the analgesic effects of TENS on various chronic pain patients, aiming to understand its clinical mechanisms better.

## Contributors

Z.W., L.H., and Y.K. designed the study; Z.W., Y.D., Y.Z., Y.L., and Y.K. performed data acquisition; Z.W., X.L., L.Z, J.C.W.B., and Y.K. performed data analysis. The original draft was written by Z.W., with Y.K., L.H., M.Z., and J.C.W.B. performing review and editing. All authors reviewed and accepted the final draft of the manuscript.

## Data sharing

Authors will share analytic code and raw, de-identified individual participant brain imaging data on request with researchers who provide a methodologically sound proposal and can conduct analyses that achieve the aims of the proposal. We are preparing a data sharing platform and associated study protocols and code, which should be completed approximately 3 months post publication. Data sharing requests can be directed to. To gain access data requestors will need to sign a data access agreement.

## Acknowledgments statement

Thanks to Jiyuan Wang, Guangyue Tian, Jiahui Zhong, and others for their support in data collection and methodology. Thanks to Siemens Health engineers who assisted in sequence commissioning and the engineers who repaired the experimental equipment.

